# Fate trajectories of CD8^+^ T cells in chronic LCMV infection

**DOI:** 10.1101/2020.12.22.423929

**Authors:** Dario Cerletti, Ioana Sandu, Revant Gupta, Annette Oxenius, Manfred Claassen

**Affiliations:** Institute of Molecular Systems Biology, ETH Zurich, Otto-Stern-Weg 3, 8093 Zürich, Switzerland; Institute of Microbiology, ETH Zurich, Vladimir-Prelog-Weg 4, 8093 Zürich, Switzerland; Internal Medicine I, University Hospital Tübingen, Faculty of Medicine, University of Tübingen, Otfried-Müller-Straße 10, 72076 Tübingen, Germany

**Keywords:** LCMV, chronic infection, single cell, transcriptome, trajectory, computational

## Abstract

In chronic infections CD8^+^ T cells acquire a state termed “exhaustion” which is characterized by impaired effector functions and expression of co-inhibitory receptors as response to continuous TCR stimulation. Recently, the pool of exhausted T cells has been shown to harbor multiple functionally distinct populations with memory-like and effector-like features, though differentiation and lineage relations between these are unclear. In this work we present a comprehensive scRNAseq time-series analysis from beginning of infection to established exhaustion in CD8 T cells. We apply lineage inference using informed cell transitions derived from RNA velocity to identify differential start and end states and connections between them. We identify a branch region early during chronic infection where pre-committed cells separate into an exhausted and a memory-like lineage and discovered molecular markers demarcating this branch event. Adoptive transfer experiments confirmed fate-commitment of cells only after this branch point. We additionally linked the progression along developmental lineages to antigenic TCR stimulation.

## 1 Introduction

Viral infections with human immunodeficiency virus (HIV), hepatitis C virus (HCV) and in mice with lymphocytic choriomenengitis virus can result in chronic infection with ongoing viral replication and high antigenic load for weeks or months. This continuous exposure to antigen drives CD8^+^ T cells into a functionally distinct phenotype, termed exhaustion [1]. This state is characterized by functional, transcriptional and epigenetic changes that result in expression of co-inhibitory receptors such as PD-1, LAG-3, 2B4 and CD160, decreased secretion of cytokines like INF*γ* and TNF*α* as well as reduced proliferation and survival. Acquisition of this exhausted phenotype is a continuous and gradual process driven by excessive TCR stimulation [2].

In this context of chronic antigen exposure, CD8 T cells undergo a differentiation program that differs markedly from the one observed during acute resolved infection. Previous studies have analyzed and inferred differentiation trajectories of virus-specific CD8 T cells using bulk or single cell transcriptomic profiling in various systems, including the model of chronic LCMV infection [3, 4, 5]. Most of these studies have inferred differentiation trajectories based on bulk or single cell analyses performed at one or two time points during chronic LCMV infection [4, 5].

Asynchronicity in this process as well as different micro-environments that CD8^+^ T cells experience result in a heterogeneous population of cells at a given time point of the infection. One sub-population of virus-specific T cells acquires a phenotype that shares properties with memory T cells from acute infection and has been linked to the expression and activity of T cell Factor 1 (TCF1) [3] [6]. In contrast to terminally exhausted or effector T cells, these cells retain proliferative activity and have better survival in the infected host [4]. It is not yet fully understood how and when these different cell states arise during the course of the infection and which intermediate cell states precede these.

Recent advances in sequencing technologies have made it possible to profile cells genome wide on the transcriptional level using single-cell RNA sequencing (scRNAseq). This technology allows capturing the transcriptional heterogeneity of multiple cell populations and to computationally infer sequences of cell states traversed during dynamic processes such as T cell differentiation in chronic infections. When analyzing scRNAseq datasets, cells are treated as points in transcriptome space based on their expression profile. Dimensionality reduction techniques like t-SNE [7] and UMAP [8] construct two-dimensional representations for analysis and interpretation of the high dimensional single-cell expression data. Pseudotime and lineage inference methods aim at constructing likely transitions between cell states [9].

Recent studies aimed at reconstructing cell state sequences of CD8^+^ T cell differentiation in chronic LCMV infection [4], [5], [10]. They discovered multiple phenotypic subsets, namely memory-like, terminally exhausted and effector-like cells and investigated likely transitions between these subsets. However, these studies lack temporal resolution to reliably infer trajectories and to identify potential branching events in the differentiation process. Samples were either generated from different infection settings at single time-points, or at far spaced time-points. Further, applied trajectory inference methods infer pseudotime and lineages based on similarity of transcripts and lack taking advantage of all the information present in the scRNA seq data. Directionality information is now – in principle – available for trajectory inference via RNA velocity analysis. RNA velocity [11] considers additional information about the ratio of un-spliced to spliced mRNA in transcript data, which serves as a measure to determine the stage (early, intermediate, late) of individual gene expressions and allows to predict the future expression state and hence to better infer the directionality towards their neighbors in the high-dimensional transcriptional space. So far no study leveraged RNA velocity in order to include this information to disambiguate the results from conventional trajectory inference.

In this work we conducted scRNAseq measurements at multiple time-points ranging from the beginning of chronic LCMV infection until manifestation of exhaustion three weeks after infection. This level of time resolution allows more detailed identification of cell states and their differentiation. We further included information from RNA velocity analysis to perform simulation based trajectory inference of differentation events leading to the different terminal CD8^+^ T cell states observed in chronic LCMV infection. This is the first attempt to make use of RNA velocity to produce informed differentiation trajectories that connect the different cell states. This analysis allowed us to construct faithful lineage trajectories towards the two endpoints of differentiation, namely a terminally exhausted and a TCF1^+^ cell population. We identified a potential branching point in the initially shared trajectories and validated our findings using adoptive transfer experiments of cells arising before or after the branching point. We confirmed that cells before the branch point gave rise to both exhausted and TCF1^+^ cells, whereas exhausted cells after the branch point maintained their phenotype. Additionally, we demonstrated that TCF1^+^ cells largely retained their phenotype in absence of antigen stimulation, corroborating the end-point differentiation characteristics of this population. However, if exposed to antigen stimulus, the TCF1^+^ population has the ability to differentiate into terminally exhausted cells, in line with previous adoptive transfer experiments.

## 2 Results

We first investigated the differentiation landscape of CD8 T cells, followed by RNA velocity analysis to reveal developmental endpoints. Afterwards we identified two branching trajectories towards memory-like and exhausted cell states, respectively. We validated commitment to the branches using adaptive transfer experiments and additionally highlight the importance of antigen stimulation during development.

### 2.1 Differentation landscape of CD8 T cells during chronic LCMV infection

We acquired single cell transcriptomic data from multiple time points during chronic infection, covering the very early phases (day 1-4), peak phase (day 7), contraction phase (day 14) and late phase (day 21) (Fig. 1), with the aim to capture an increased spectrum of the transcriptional landscape during the course of the infection that would allow a time-resolved analysis of single cell heterogeneity and possibly more accurate inference of differentiation trajectories of virus-specific CD8 T cells. To this end, T cell receptor (TCR) transgenic (tg) LCMV gp33-41-specific CD8 T cells (P14 cells) were adoptively transferred into naïve C57BL/6 mice, followed by infection with LCMV clone 13 (Cl13). Activated and expanded P14 cells were isolated at the above indicated time points and subjected to single cell RNAseq (scRNAseq) analysis using the 10x Genomics platform.

**Figure 1:**
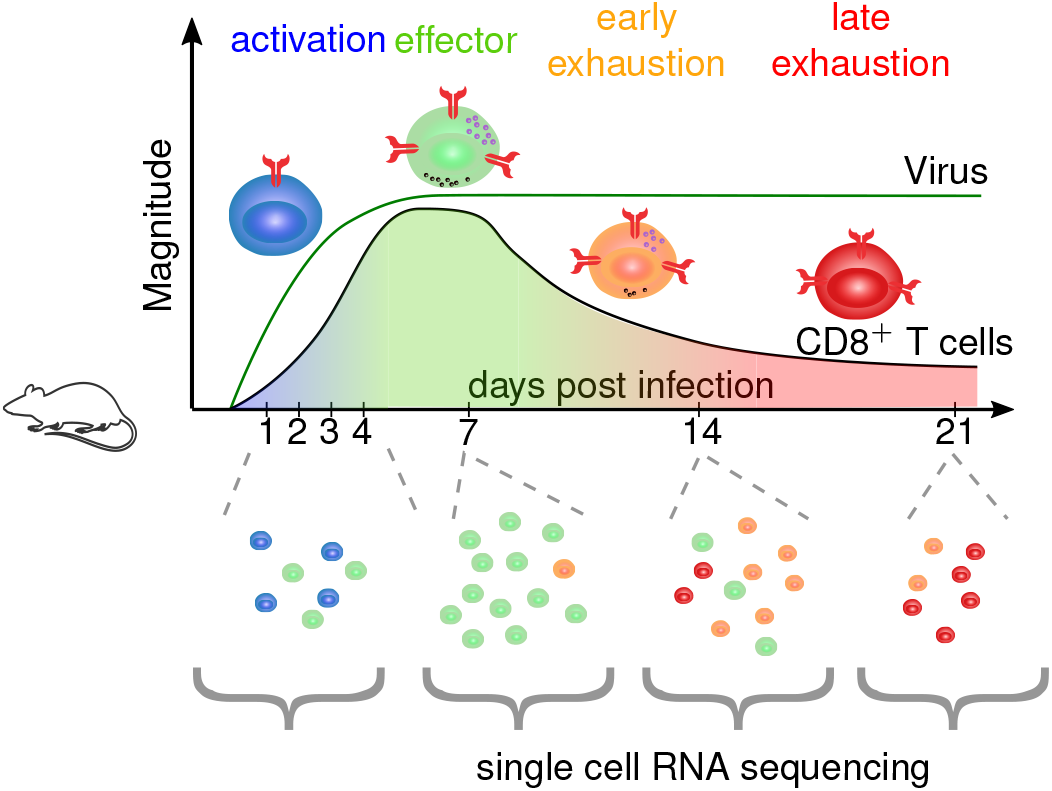
Transgenic P14 CD8 T cells were sampled longitudinally during infection. The samples were acquired from four phases of the infection activation (day 1-4), effector (day 7), early exhaustion (day 14) and late exhaustion (d21) and scRNAseq was performed using the 10x Genomics platform.

For exploratory analysis of the transcriptome data of the multiple time points, we applied commonly used filtering and scaling of the raw data [12] and applied principal component analysis to reduce noisy signals. The resulting multidimensional data was projected into two dimensions using UMAP [8].

The UMAP projection revealed a continuous structure of the data emerging from the time-resolved samples, supporting a continuous developmental process (Fig. 2a). Louvain clustering [13] defined 11 distinct clusters across all samples (Fig. 2b,d, Supp. Fig. S3). Based on previously described markers for the exhausted subsets, such as CD160, CX3CR1 and TCF1 [3, 1, 4], we further aggregated these cluster into 6 phenotypic groups (Fig. 2c). Activated cells from day 1 to 4 post infection (dpi) clustered at one peripheral region in the data, termed in the following **early group** (green in Fig. 2c). Differential expression revealed that the early group presented expression patterns of proliferation as well as of exhaustion, indicated by expression of *Mki67, Cdca3* but also *Cd160*.

**Figure 2:**
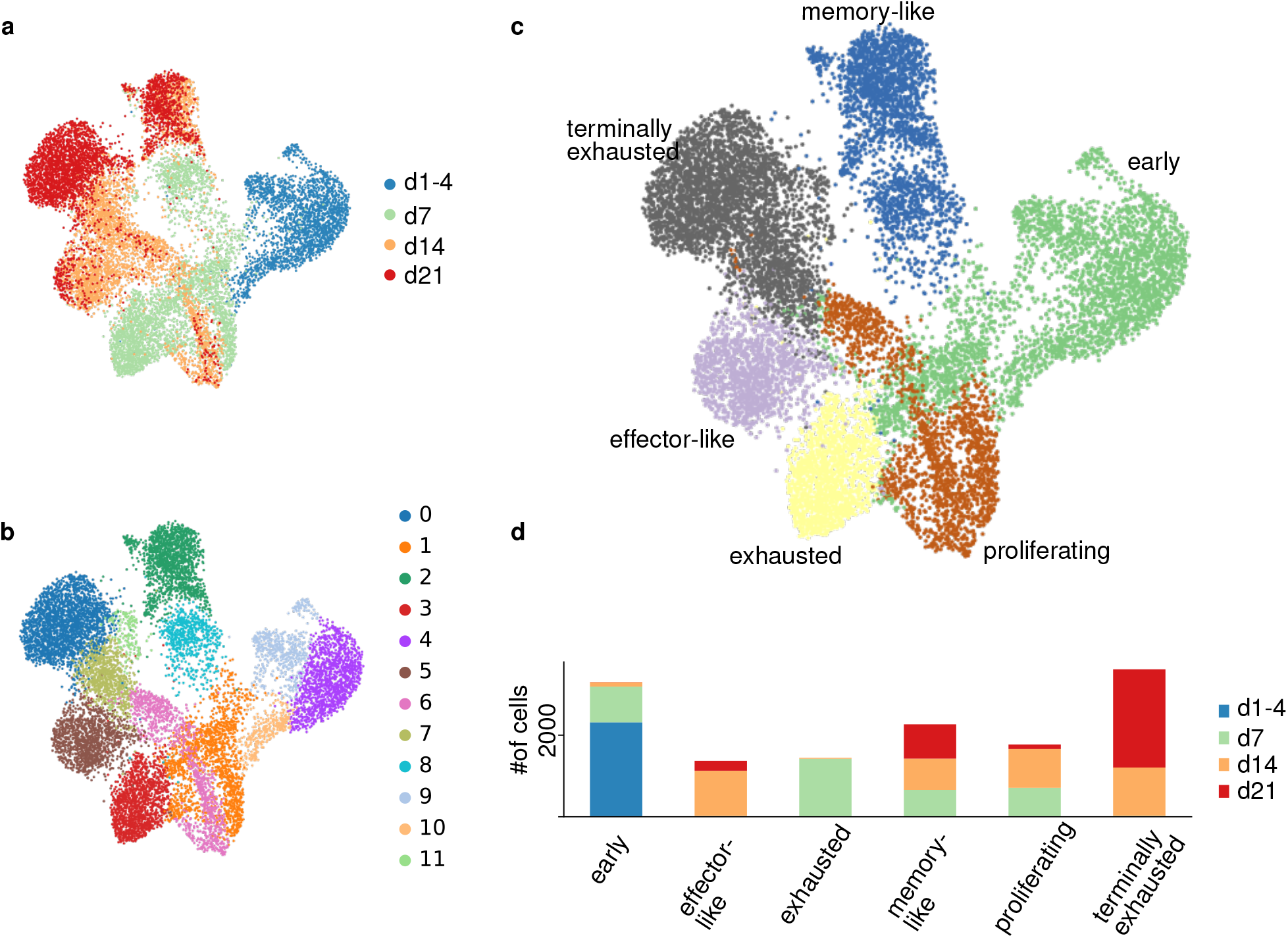
UMAP projections of the filtered and normalized transcript counts are shown. (a) Color indicates the time-point after infection, at which cells were isolated for scRNAseq (b) louvain cluster assignment based on the first 50 PCs (c) phenotypic cluster annotation based on previous marker genes and differentially expressed genes [4] (d) Cell composition of the phenotypic clusters by sample time-point.

Conversely, two distinct populations from the latest time point at 21 dpi clustered at the other extremes of the spectrum. Of these “late” endpoints, one population showed high expression of a number of inhibitory receptors, including *PD-1 (Pdcd1), CD39 (Entpd1), LAG-3 (Lag3)* and *CD160 (Cd160)* (Fig. 3a, b), indicating a terminally exhausted phenotype [14]. This **terminally exhausted group** (grey in Fig 2c) was composed of clusters from d7 and 14 and had the highest expression in co-inhibitory receptors and additionally showed high expression of the transcription factor *EOMES*. The other end-point populations showed high expression of the transcription factor *Tcf7* (Fig. 3a, b), the memory-marker *Il7r* as well as *Slamf6*, revealing this cluster as the previously described memory-like population [3]. This **memory-like group** (blue in Fig 2c) was composed of two clusters from day 7, 14 and 21, all having high expression of *Tcf7*.

**Figure 3:**
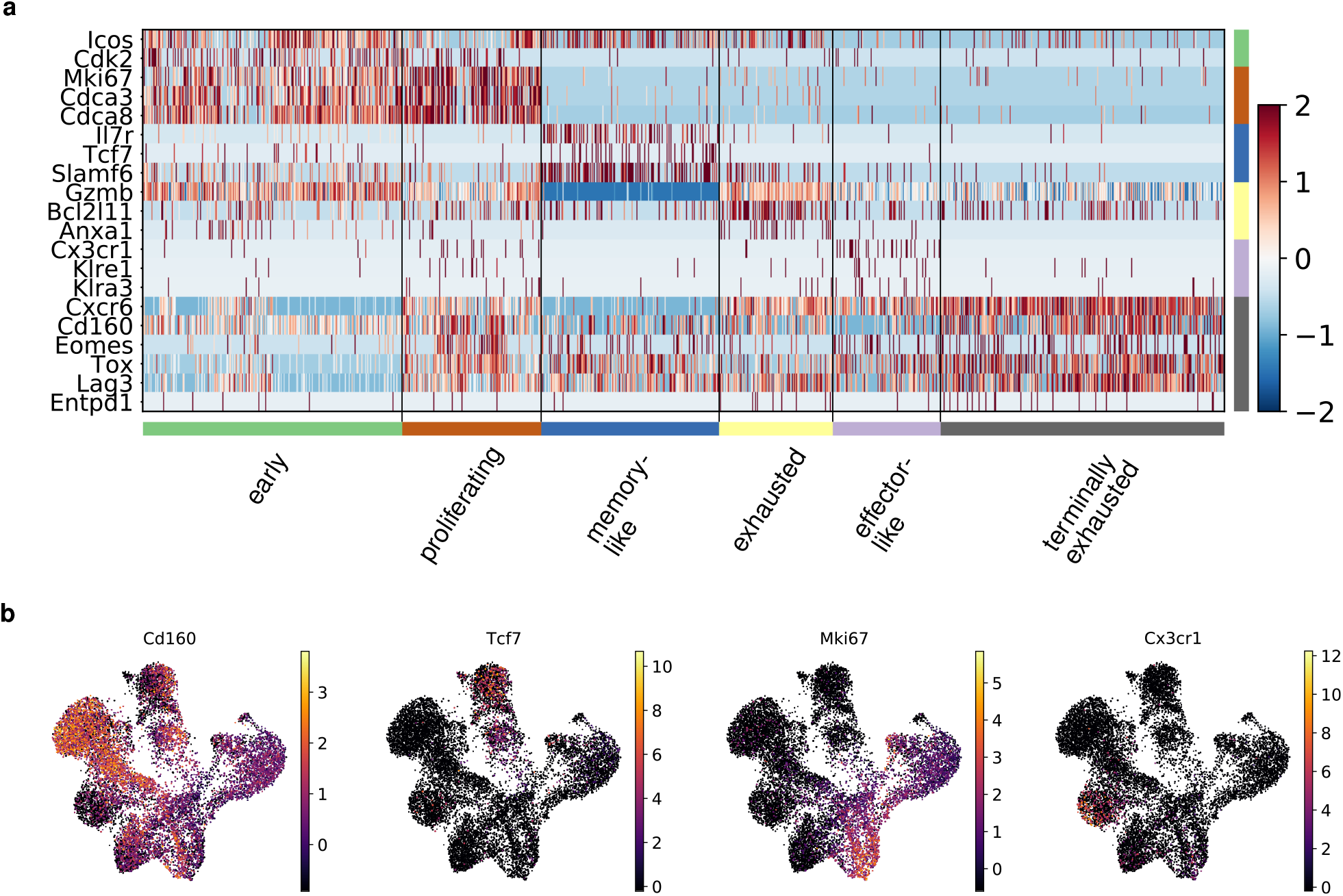
(a) Heatmap of normalized gene expression for a selection of group specific genes. Columns are individual cells arranged by phenotypic groups. The genes in the rows are grouped according to their phenotypic assignment. (b) UMAP projection of expression pattern of identified group specific genes for the terminally exhausted (Cd160), memory-like (Tcf7), proliferating (Mki67) and effector-like group (Cx3cr1)

Cells from 7 and 14 dpi were situated in between the 14 dpi and 21 dpi samples, with the 7 dpi samples being similar to cells from early time-points on one end of the spectrum, but also connecting to already splitting trajectories into exhausted and memory-like populations on dpi 14. At day 7 was one cluster identified that presented clear signatures of exhausted CD8 T cells but retained some expression of *Gzmb* but also apoptotic genes like *Anxa1*, we termed this the **exhauted group** (yellow in Fig. 2c).

At 14 dpi, some of the cells were still connected to the 7dpi cell states but a considerable fraction of cells had already further differentiated towards the two endpoints of 21dpi, in particular towards the memory-like endpoint. One cluster expressing *Cx3cr1* exclusively was termed **effector-like group** (purple in Fig. 2c). Differential expression analysis (Fig. 3a, Supp. Fig. S3) revealed higher expression levels of *Cx3cr1* and additionally killer lectin receptor genes (*Klre1, Klra3*). These effector-like cells were only present in samples from day 14 and 21.

We also identified a strong cell cycle component in the two clusters presenting high expression levels of e.g. *Mki67* (cluster 1 & 6) (Fig. 3b). Further, we calculated scores for cell cycle and cell division genes based on the three different cell cycle stages G1 phase, S phase and G2/M phase (Supp. Fig. S4). We observed that this **proliferating group** (brown in Fig. 2c) presented high scores for G2/M phase and was composed of cells from the 7 and 14 dpi and to a lesser degree from the 21 dpi time-point.

### 2.2 RNA velocity analysis reveals developmental end points

Having mapped serial single cell transcriptomes into a continuous landscape of cellular states, we next aimed at inferring informed differentiation trajectories into this landscape. To this end we leveraged RNA velocity [11] and applied it to our longitudinal data set, revealing a vector field demarcating likely transitions between all cell states in the dataset. Applied to our dataset, this analysis clearly revealed a transition flow from early activated cells at 1-4 dpi towards early (dpi 7) and late exhausted cells (dpi 14 & 21) (Fig. 4a).

**Figure 4:**
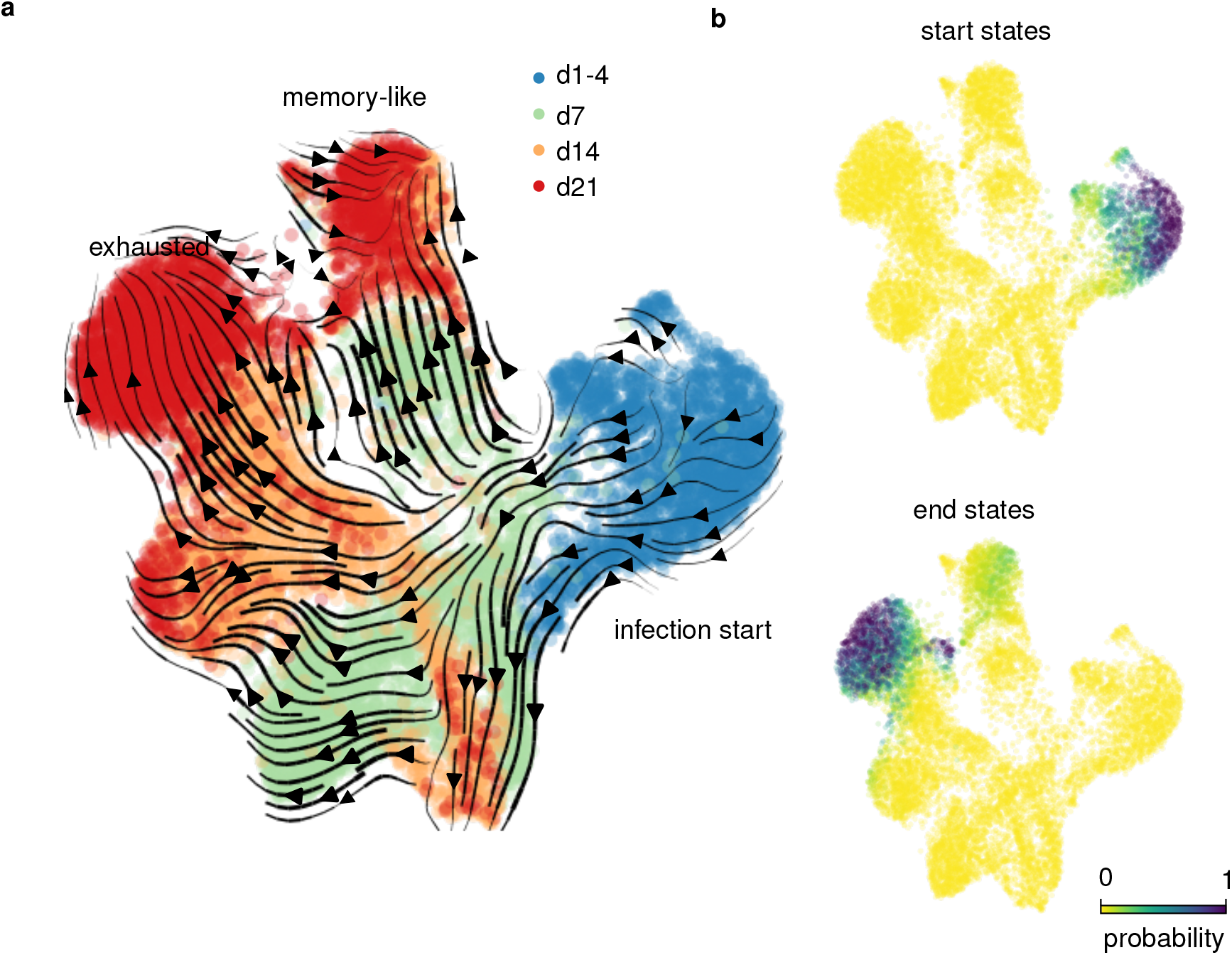
(a) stream plot visualizing likely transitions between cells inferred from RNA velocity (b) The stationary distribution of the backward and the forward transition matrix, respectively, indicate start and end cell states.

**Figure 5:**
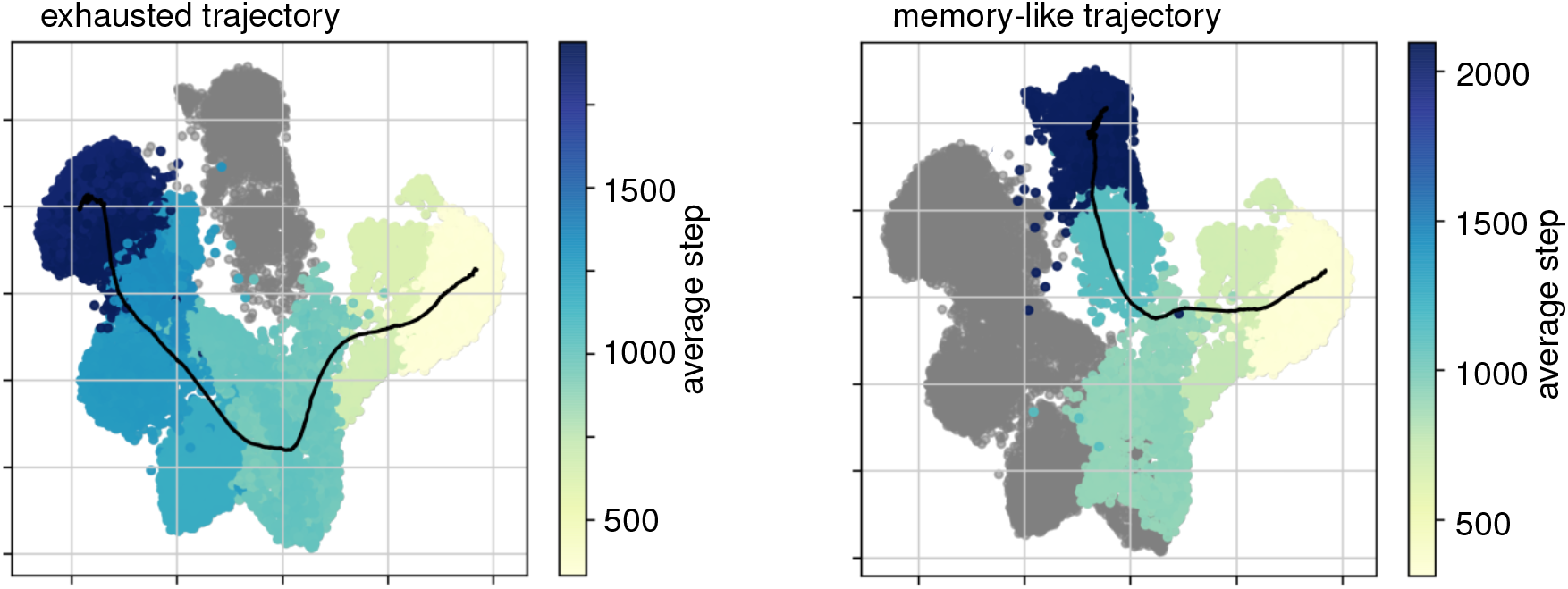
Lineage trajectories after simulations towards the two endpoint populations using cytopath. The average simulation steps to arrive at a cell are color coded per cluster. The coordinates of the average trajectory after alignment is depicted as black line. The shown average trajectories are based on 2000 simulations per endpoint.

Using the calculated cell transitions, we defined a Markov process with cells as the states and transition probability estimates from RNA velocity (see Method section). Computing the equilibrium distribution of the forward Markov process corresponded biologically to the most differentiated phenotypes as the cells transitioned to more differentiated states until they had acquired their final transcriptional state and did not differentiate further. Conversely, we inverted the transition probabilities and computed the equilibrium of the backward Markov process. Cell states thereby transitioned to their most likely previous transcriptional state resulting in the most undifferentiated state, having the highest probability.

This allowed us to assign cells that are most likely at the start and at the end of the differentiation process. The highest probably of a start region was at the edge region of the early group (Fig. 4b). This seemed plausible, as we would expect that the differentiation process started at the edge of the earliest sample (dpi 1-4), where there are no preceding cell states. This region showed gene signatures indicating strong DNA synthesis and cell cycle activity, that conferred an activated phenotype (Supp. Fig. S6).

The highest probability for end points was in regions from dpi 21 in the terminally exhausted group (0.5 on average in cluster, maximum 1.0). In this region many signaling related genes had changed expression, which could be a result of co-inhibitory receptor signaling (Supp. Fig. S6). Additionally, there was a local maximum in end point probability in the memory-like group from dpi 21 (0.1 on average, maximum 0.3). Differentially expressed genes in this region comprised typical genes of the memory-like signature, namely *Il7r* and *Tcf7* (Supp. Fig. S6). Both the terminally exhausted cells as well as the memory-like cells seemed to comprise an endpoint of differentiation state.

We assessed and confirmed the robustness of the velocity fields by confirming the practical equivalence of start and endpoint estimates across 574 parameter variants (Supp Fig. S5).

RNA velocity analysis additionally indicated that the process from the start to the end-points is gradual, since there were a multitude of intermediate transcriptional states between the two extremes. We observed also transitions between all these intermediate states (Fig. 4a).

### 2.3 Simulation based inference reveals trajectories towards exhausted and memory-like phenotypes

Based on the high-dimensional vector field resulting from the RNA velocity analysis, we calculated a transition matrix that contains likely future states for each cell. This transition matrix we used to simulate differentiation trajectories from cells in the start region. We aimed at understanding the developmental paths that an activated CD8^+^ T cell could follow to acquire the two differentiated end-point phenotypes. The probability of a cell moving from one transcriptional state to another can be approximated by the transition probabilities from RNA velocity. We used the calculated root cells (Fig. 4b) as starting points for stochastic simulations. Each differentiation step in the transition matrix was simulated according to the transition probability until one of the previously defined end stages (Fig. 4b) was reached. This sequence of steps approximated one possible path of differentiation for each cell (Supp. Fig. S7). We simulated 2000 trajectories per endpoint to sample the whole spectrum of possible differentiation trajectories. We observed a strong disbalance in preference for the endpoints. Only about 1% of the simulated sequences ended up in the memory-like cluster, whereas the remaining ones differentiated into the exhausted endpoint. We expected the ratio to be shifted towards the exhausted phenotype but not to this extent, since we measured around 10% memory-like cells at day 21. We performed more simulations towards the memory-like endpoint to balance the number of trajectories. All the obtained trajectories were then aligned using dynamic time warping and clustered to generate average trajectories. Each cell was assigned to the nearest average trajectory according to the alignment score (Method section 4), calculated from cosine distance of its RNA velocity with the direction of the trajectory (Fig. 6a). This resulted in a temporal ordering of the cells in conjunction with a score to which trajectory it belongs. The detailed procedure of Cytopath is described in [15]).

**Figure 6:**
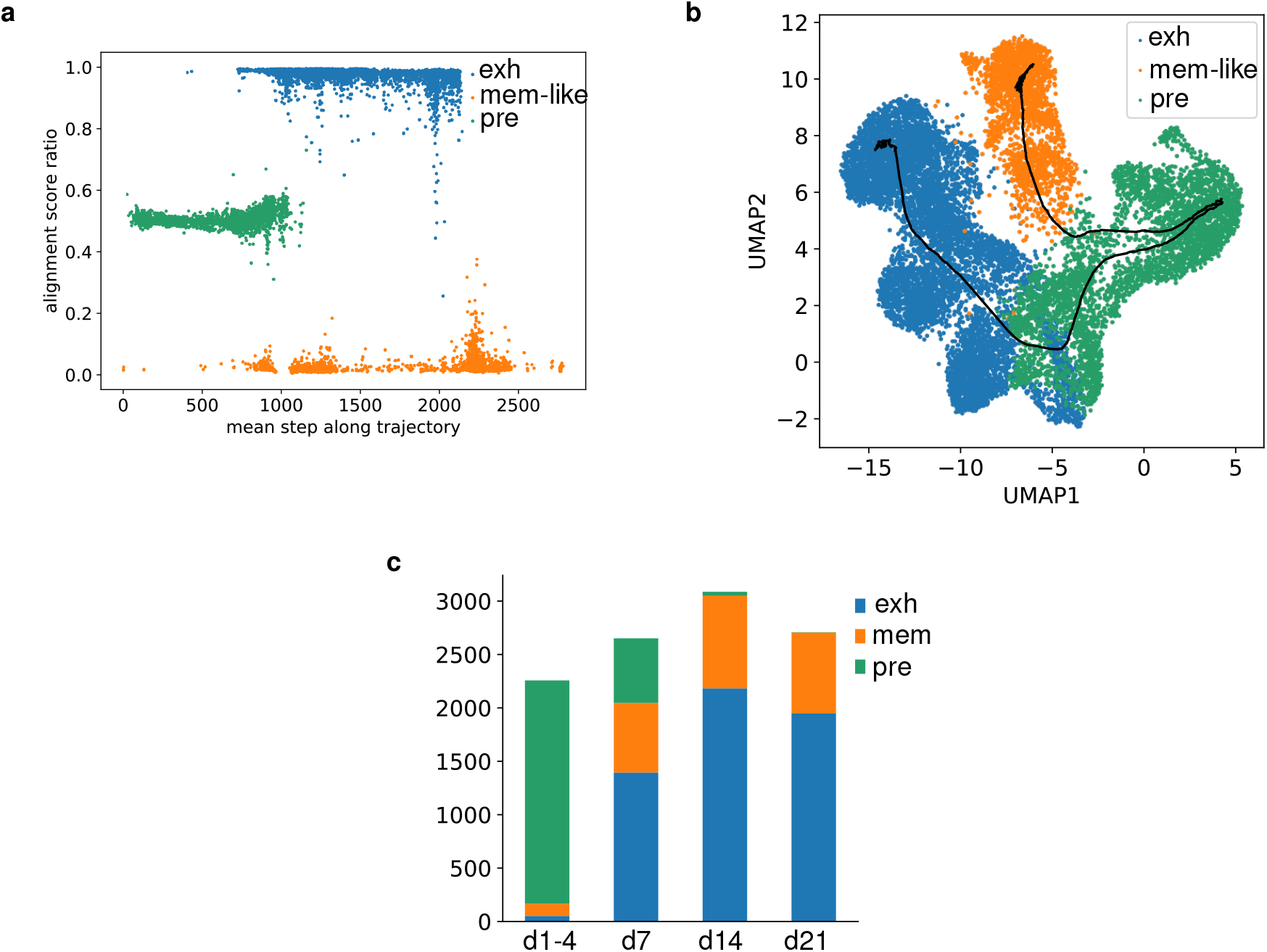
Cells are aligned to the branches of the two fates exhausted (exh), memory-like (mem) or the pre-commited branch (pre). (a) The two average trajectories (black) towards the exhausted (blue) and the memory-like fate (orange) with an pre-commited shared part (green). (b) Alignment score ratio along the cytopath simulation steps. High values indicated preferential alignment with the exhaustion trajectory, low values alignment with the memory-like trajectory. (c) sample time-point composition with respect to branch assignment.

Our analysis revealed two main trajectories, one towards the exhausted endpoint the other to the memory-like endpoint (Fig. 5). Both trajectories shared the same cell populations up to a region that was composed of cells obtained from around 4 dpi. From thereon the trajectories started to diverge into the two phenotypic branches. The differentiation trajectory towards the exhausted path included the cell population with high cell cycle activity. The memory-like trajectory did not seem to pass the region of high proliferation, but diverged earlier and transitioned towards the memory-like endpoint.

Multiple genes were found to be differentially expressed between the two trajectories (Supp. Fig. S8). In the memory-like trajectory *Slamf6, Ccr6, Tnfsf8, Xcl1* and *Cxcl10* were expressed at higher levels. Many of them showing gradually increasing expression towards the differentiated endpoint (*Slamf6, Ccr6, Tnfsf8*). *Xcl1* was highest at the start of the trajectory and later decreased but was still maintained at much higher levels than on the exhausted branch. The exhausted branch showed increasingly higher expression of *Cxcr6, Ccl5* and *Nkg7* as the trajectory progressed towards the end point. The two genes *Ifngr1* and *Lgals3* were transiently upregulated in the exhausted trajectory exclusively, but decreased towards the end point (Supp. Fig. S8).

### 2.4 Branching point of differentiation towards the end points

Based on the simulated mean trajectories we wanted to identify a branching point at which the trajectories of cells moving towards the exhausted and the memory-like endpoints would diverge. This branching region would demarcate the point after which a cell would be committed to only one endpoint.

We therefore computed the ratio of the alignment score for each cell to the two average trajectories (Fig 6a). We observed that in the unstructured early region all cells had equal scores for both the exhausted and the memory-like fate. However, between average simulation step 800 and 1200, the cells started to be uniquely assigned to only one of the two trajectories. Interestingly, we observed a region along the differentiation trajectories where some cells aligned clearly to the exhausted trajectory, some to the memory-like trajectory and some to both. It seemed that this region was where bifurcation took place. Using a threshold on the alignment score we assigned all cells to either the exhausted branch (blue), memory-like branch (orange) or pre-committed branch (green, Fig. 6).

To determine which biological time-point would correspond to the identified branch region, we investigated the branch composition of the four samples (Fig. 6c). The earliest samples were almost exclusively composed of pre-committed cells, whereas the day 7 sample contained already a large fraction of lineage-committed cells. We reasoned that bifurcation must take place between day 5 and 6 after infection.

Differential gene expression between the branching region and its immediately adjacent committed branches did not reveal any clear transcriptional signatures, that would precede or succeed bifurcation. However the trajectory assignment clearly implied that after the branching region, cells were fully committed to their lineage. Since the alignment score also considers the velocity direction of each cell, alignment to only one trajectory indicates that differentiation will take place along this path. The abrupt increase in the alignment score after the branch region suggests that cells beyond the bifurcation do exhibit coherent and significant velocity away from the bifurcation and this process is unlikely to be reversible.

### 2.5 Identification of marker genes predictive for different developmental fates

To further demarcate markers that would specify the branching point, we searched for genes that were characteristic for this bifurcation - either being expressed at divergent levels before, at, or after the branching point. We trained a classifier to predict the assigned branch label from the transcriptional profile of each cell (see Method section). If a gene was expressed in one branch but not the other, it was considered relevant for the prediction. To validate the identified branching markers later on a protein level using flow cytometry, we restricted the transcriptional input to genes transcribing proteins with validated antibodies for staining. The result of this analysis variant allowed us to sort and experimentally analyze the differentiation potential of pre-committed and committed cell states in later validation experiments.

The classifier identified 12 genes that were most relevant to distinguish the three branches (Fig. 7). These included already described markers of the memory-like population, such as *Il7r* (IL7R), *Tcf7* (TCF1) and *Slamf6* (Ly108) [3, 4], but revealed also potential new candidates *Icos* (CD278), *Ly6e* (SCA-2) and *Itgb1* (CD29) that are highly expressed in cells from the memory-like branch. Markers relevant for the exhausted branch contained *Cxcr6* (CXCR6), *Ifngr1* (IFNGR1) and *Cd3g* (CD3G) but also *Selplg* (CD162), of which only CXCR6 has been linked previously to exhaustion [16]. The pre-committed branch showed high expression of *Gzmb* (Granzyme B) and Mif (MIF) both of which were expressed at lower levels in the other lineages.

**Figure 7:**
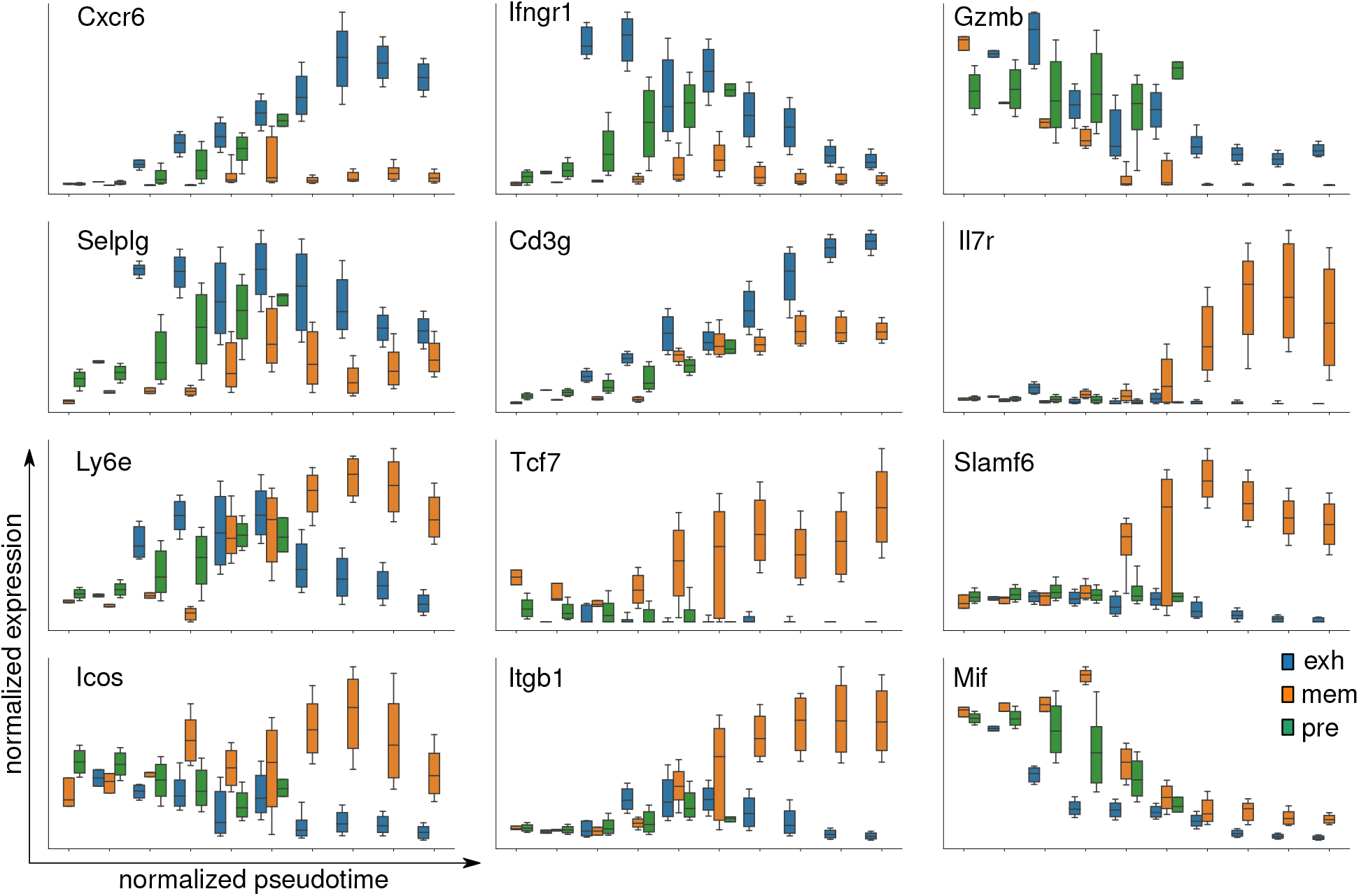
The 12 genes identified by the classifier to predict the three branch labels. Boxplot of normalized expression are shown along normalized pseudotime from cytopath. Color indicates the branch the cells were assigned to based on the Cytopath alignment score, exhausted (exh), memory-like (mem) or the pre-commited branch (pre).

Predicting the branch labels using only these markers resulted in good prediction accuracy (0.85). Additionally, using only these genes as input to UMAP showed a good separation into the three branches (Supp. Fig. S9).

### 2.6 Experimental transfer of cells with a presumably pre-committed or committed phenotype show distinct differentiation potential

To experimentally validate our branch classification, we set out to identify and later sort LCMV-specific CD8 T cells with phenotypes indicative of a pre-committed state, a committed state towards the exhausted endpoint or the memory-like endpoint. For validation of phenotypic markers classifying these three cell states, we tested all 12 identified markers for their ability to discriminate early populations to then test their fate potential. Specifically, we transferred naïve P14 cells into naïve C57BL/6 mice, followed by chronic LCMV infection. 5 days post infection (when representatives of all three populations of interest had formed, i.e. the uncommitted cells and the committed exhausted and memory-like cells), P14 cells were analyzed according to the markers identified using the classifier (data not shown). We first identified protein markers that showed high variance at the branching time-point. We determined CXCR6 and TCF1 as the prime candidates for sorting the branches into pre-committed (CXCR6^*−*^ TCF1^*−*^), memory-like (CXCR6^*−*^ TCF1^+^) and exhausted (CXCR6^+^ TCF1^*−*^) cells.

Having identified markers that allowed to distinguish between uncommitted and committed cells into the exhausted and memory-like branch, we used these markers to isolate the respective populations 5 days into chronic LCMV infection. Specifically, we transferred naïve P14 T cells expressing GFP under the TCF1 promoter into C57BL/6 mice and infected them with high dose LCMV Clone-13 (Fig. 8a). At dpi 5 we sorted P14 cells from the three branches according to expression of CXCR6 and TCF1 (detected by GFP) and transferred them into infection-matched hosts (Fig. 8b). At one week after transfer (at dpi 12 from the initial infection), we analyzed the progeny of cells originating from the three branches in the spleen (Fig. 8c). We observed that cells recovered after transfer of exhausted cells into infection-matched recipients retained their exhausted phenotype. Cells recovered after transfer of the memory-like branch exhibited phenotypes of both exhausted and memory-like cells, confirming previous results of differentiation from memory-like into exhausted cells [4] but contradicting our finding of a memory-like endpoint. Recovered cells after transfer of pre-committed cells exhibited both a memory-like or an exhausted phenotype, confirming their differentiation potential into both memory-like and exhausted cells. However, there was a strong bias towards differentiation along the exhaustion branch, which might be explained by much more extensive proliferation of these cells compared to memory-like cells.

**Figure 8:**
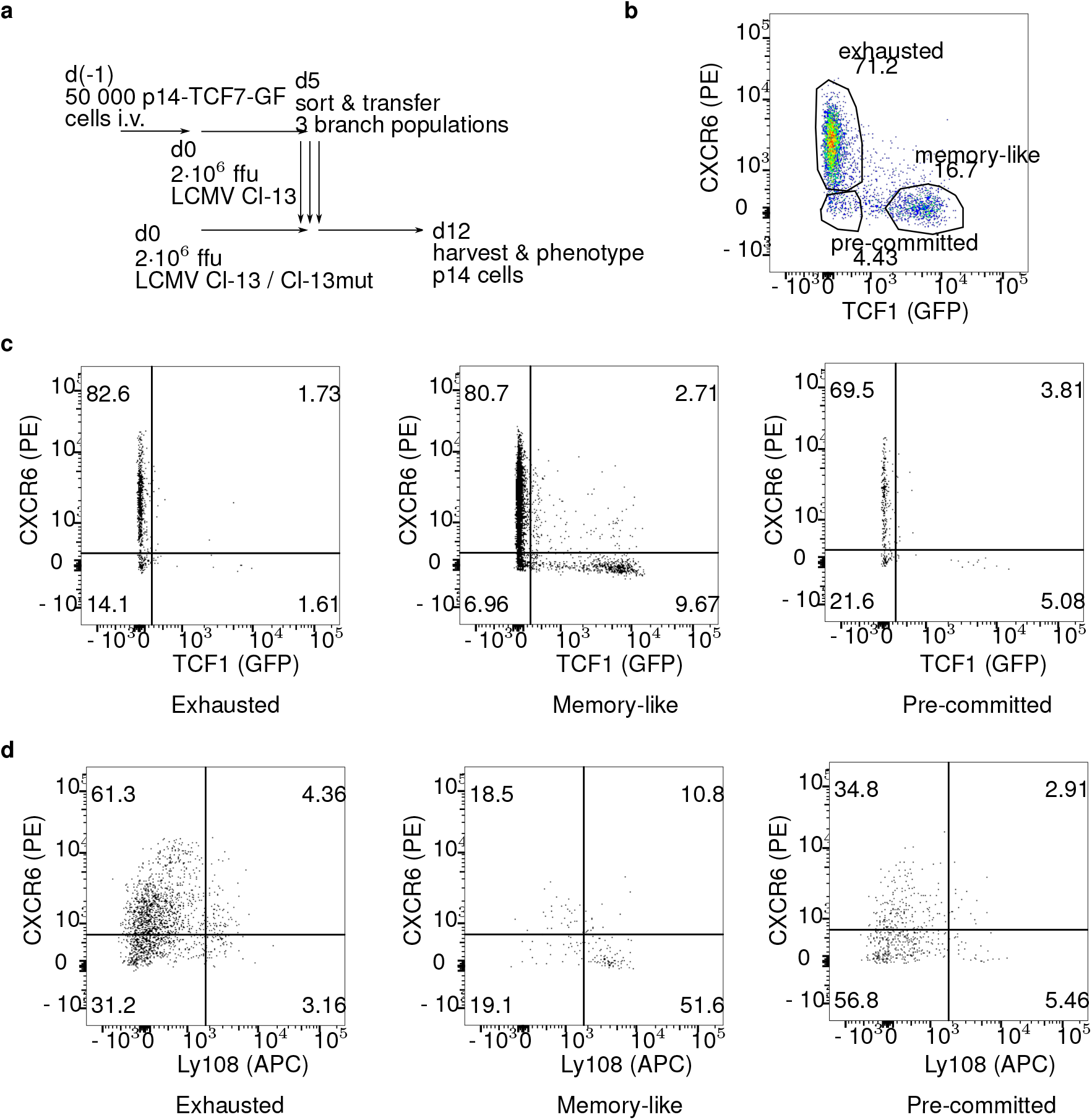
(a) The three P14 branch populations were isolated at 5 dpi from high-dose Clone-13 infected mice (that had been transferred with naive P14 cells prior to infection) and transferred into infection matched hosts and their phenotype was assessed 7 days post transfer. (b) flow cytometry gates used to sort exhausted, memory-like and pre-committed branch (c) phenotype of the recovered P14 cells at 12 dpi from spleens after high-dose Clone-13 infection and transfer of either exhausted, memory-like or pre-committed cell populations isolated at 5 dpi. Cells are gated on P14 cells. (d) phenotype of the recovered cells at 12 dpi from spleens after transfer into hosts infected with Clone-13 P14 escape mutant. Naïve P14 cells were first transferred into naive C57BL6 mice, followed by Clone-13 infection. At 5 dpi, exhausted, memory-like and pre-committed populations were sorted and adoptively transferred into infection matched hosts with Clone-13 escape mutant. Recovered P14 cells are shown.

### 2.7 Differentiation transitions are driven by antigenic TCR stimulation

Since our RNA velocity and trajectory analysis had revealed the memory-like cells as an end-point of differentiation, we speculated that their unexpected differentiation into terminally exhausted cell states after adoptive transfer is triggered by external cues that drastically changed their state. We investigated the possibility of TCR stimulation as such a cue by transferring cells belonging to the three branches (isolated from mice at 5 dpi following adoptive transfer of P14 cells and induction of chronic infection with Clone-13) into hosts infected with a LCMV Clone-13 strain that expresses a variant of the gp33 peptide that is not recognized by P14 cells (Fig. 8a).

We recovered cells from spleen and lymph nodes and analyzed their phenotype based on CXCR6 and Ly108 expression (Fig. 8d). The recovered cells after transfer of the exhausted branch again retained their exhausted phenotype. Surprisingly, we recovered much fewer cells with an exhausted phenotype after transfer of memory-like cells and a major fraction retained their memory-like phenotype, indicating that further differentation was largely halted in the absence of antigen. The transfer of pre-committed cells resulted in recovery of cells with a largely undifferentiated phenotype of neither terminally exhausted nor memory-like, largely retaining their pre-committed state. These results clearly pointed towards TCR stimulation being a major driver of differentiation during chronic infection for both the pre-committed state and for further differentiation of the memory-like state into fully exhausted cells.

## 3 Discussion

We analyzed differentiation trajectories of virus-specific CD8^+^ T cells during chronic LCMV infection using scRNAseq time-series data from four different time points covering activation, peak, contraction and late phase of the response. The time-resolved traversal of the transcriptional landscape revealed a continuous and bifurcating process, with early activated cells at the beginning and both terminally exhausted as well as memory-like cells at the end of this process. We observed cells with an exhausted phenotype and an effector-like cell population as transient states during early and late stages of this process, respectively. We further observed for all time points a population of cells with transcriptional profiles of proliferation.

Applying RNA velocity analysis to our single-cell transcriptional data allowed us to estimate transitions between the cell states during the progressing immune response. We computed most likely end and start regions and identified two major differentiation paths leading to the exhausted population and the memory-like population respectively. At the early time-points until about day 5, the two average trajectories were nearly indistinguishable, but then diverged increasingly towards their respective endpoints.

We identified a branching region in the early stages of infection before day 5 post infection. Before this branching point, pre-committed cells would still have the potential to differentiate into both the exhausted and the memory-like phenotype, whereas cells that had passed the branching point would be destined to differentiate into the endpoint they had committed to.

Although it is conceivable that factors that were not revealed by transcriptional analysis might be involved to already pre-determine cell fate during early activation [17] and regulate differentiation, this is not evident on a transcriptional and protein expression level, where branching of the two main fates manifested itself around day 5 post infection.

We derived a small set of gene markers that separate cells into the pre-committed (TCF1^*neg*^ CXCR6^*neg*^), exhausted (TCF1^*neg*^ CXCR6^*hi*^) and memory-like branch (TCF1^*hi*^ CXCR6^*neg*^) by a classification model. Adoptive transfer of cells sorted according to these markers, and thereby likely belonging to the three branches, at day five post chronic LCMV infection into infection-matched new hosts, confirmed the plasticity of pre-committed cells to acquire both exhausted as well as memory-like phenotype. Conversely, progeny from committed exhausted cells largely retained their phenotypes. Although, our velocity and trajectory analysis suggested the memory-like cells to represent a developmental end-point, we still recovered both memory-like and exhausted cells after transfer of cells from the memory-branch. These transitions might be rare (and fast) events in the integral setting of an infected mouse, and thus not be represented in the scRNAseq data. Additionally, i.v. adoptive transfer of memory-like cells into circulation might expose them to different antigenic burden compared to their natural niches, thereby accelerating a differentation process.

Previously published work used scRNAseq to study CD8 T cell differentiation during chronic infection, isolating virus specific T cells at different time-points [4, 5, 10, 18]. They made use of dimensionality reduction and computational tools for trajectory inference. Although, these studies used fewer time-points and the lineage inference included one time-point only. All these studies consistently described the memory-like and the terminally exhausted cell state, which we also identified. Some studies [5, 18] additionally described an effector-like CX3CR1^+^ population arising late during the infection, which we also found in our late samples from 14 & 21 dpi.

Two studies [4, 5] investigated plasticity and differentiation of the memory-like cells computationally by lineage inference on a single time-point and through adoptive transfer experiments, concluding that memory-like cells partially maintain their phenotype and can give rise terminally exhausted and effector-like cells. Our lineage inference across multiple time-points suggests that the exhausted and the memory-like lineages are separate. We could not exclude, that we missed certain cellular states in our data set, since we only studied CD8 cells isolated from the spleen of infected animals. We were unable study developmental stages that are spatially restricted to lymph nodes or specific anatomical regions within secondary lymphoid organs, or specific differentiation processes that are restricted to non-lymphoid organs [16]. Additionally, if cell state transitions are rare or fast, it is unlikely that we would capture them in our snapshot analysis.

Our adoptive transfer experiments of memory-like cells revealed extensive transitions to terminally exhausted states, which our lineage inference did not detect. However, our transfer experiments into Clone-13 P14 escape mutant infected hosts suggested that theses transitions were strictly dependent on antigenic TCR stimulation. This could imply that we did not observe these transitions in our scRNAseq data because memory-like cells receive little TCR stimulation from their microenvironment in a natural setting. Isolation of memory-like cells by removing them from their niche and transferring them via intravenous injections could expose them to excessive antigen and trigger differentiation towards terminally-exhausted cells.

Chen et al. [4] studied early bifurcation events towards either an effector state or TCF^+^ precursor state. However, the population they termed “effector” cells already expressed many co-inhibitory receptors like our exhausted group from 7 dpi. They found this early effector cells to be very short-lived and disappear between 8 and 12 dpi. Although our data suggested a similar bifurcation into effector and memory-like lineage, our exhausted trajectory placed the early exhausted cells as an intermediate state on the differentiation towards terminally exhausted states. Since Chen et al. used KLRG1 to identify their effector population, the disappearance of these cells could be explained by down-regulation of *Klrg1* during differentiation towards terminal exhaustion.

The velocity based endpoint analysis did not reveal either the early exhausted or the effector-like states to compose stable end-points, but that all those states differentiated into terminally exhausted cells. The velocity transitions did show some flow out of these populations though, which could indicate migration out of the tissue or apoptosis of these cells, although we did not find strong apoptotic signatures in our data. Considering, that apoptotic cells are cleared very fast by the phagocytes, clearance of apoptotic cells might be too fast to capture.

Our classification analysis of the branch point revealed a set of genes that discriminates between the three branches. Even though, our validation experiments confirmed that CXCR6 and TCF1 expression patterns capture the differentiation potential of CD8 T cells early during the infection, other identified genes might still be relevant in shaping this bifurcation. Both genes *Ifngr1* and *Selplg* are transiently upregulated around the birfucation point, which could imply some influence on cell fate. Considering, that withdrawing T cells from antigen stimulation practically halted differentiation of pre-committed cells and maintained their state, points towards a significant role of TCR stimulation and signaling in this bifurcation and decision process.

This work provides additional insights into the differentiation process of CD8 T cells using a combined approach of scRNAseq analysis, computational trajectory inference and adoptive transfer experiments. Our study revealed an early bifurcation event, that shaped the differentiation fate during the course of a chronic infection and additionally highlights TCR stimulation as a significant driver of this differentiation.

## 4 Material & Methods

### Infections and cell isolation

#### Mice

Wild-type male C57BL/6J mice were purchased from Janvier Elevage. Nr4a1-GFP mice expressing GFP under the control of the NUR77 promoter [19], P14 transgenic (CD45.1) mice expressing a TCR specific for LCMV peptide gp33–41 [20] and TCF1-GFP mice expressing GFP under the control of the Tcf7 promoter [3] were housed and bred under specific pathogen–free conditions at the ETH Phenomics Center Hönggerberg. All mice used in experiments were between 6–16 weeks of age. P14-Nr4a1-GFP mice were generated by crossing Nr4a1-GFP mice to P14 mice. P14-TCF1-GFP mice were generated by crossing TCF1-GFP mice to P14 mice. All animal experiments were conducted according to the Swiss federal regulations and were approved by the Cantonal Veterinary Office of Zürich (Animal experimentation permissions 147/2014, 115/2017).

#### Virus

LCMV clone 13 [21] was propagated on baby hamster kidney 21 cells. LCMV clone 13 P14 escape mutant [2] was propagated on MC57G cells. Viral titers of virus stocks were determined as described previously [22].

#### Infection

10^4^ transgenic cells (P14, P14-TCF1-GFP or P14-Nr4a1-GFP) were adoptively transferred 1 day prior LCMV clone 13 intravenous (IV) infection with 2 x 10^6^ ffu/mouse. For isolation at 1, 2, 3, 4 days post-infection, 10^5^ P14-Nr4a1-GFP cells were transferred.

#### Cell isolation from tissues

After 1, 2, 3, 4, 7, 14 and 21 days of chronic infection, mice were sacrificed with carbon dioxide and organs (spleen, lymph nodes) were isolated. Spleens and lymph nodes were mashed through 70 µm filters with a syringe (1 mL) plunger. Cell suspensions were filtered (70 µm) and treated with ammonium-chloride-potassium buffer (150 mM NH4Cl, 10 nM KHCO3, 0.1 mM EDTA in water) to lyse erythrocytes for 5 min at room temperature.

#### Cell sorting

Spleen samples were depleted of CD4 and B cells by incubating splenocyte suspensions in enrichment buffer (PBS, 1%FCS, 2 mM EDTA) with biotinylated *α*-CD4 and *α*-B220 antibodies at room temperature for 20 min, followed by incubation with streptavidin-conjugated beads (Mojo, Biolegend) (4%) for 5 min at room temperature. Cells were then placed on a magnetic separator (StemCell) for 10 min at room temperature, followed by collection of supernatant. For scRNAseq samples, cell suspensions of spleens isolated from five mice were pooled in samples from day 7, 14 and 21 post infection and from three mice for samples from early timepoints at day 1, 2, 3 and 4 order to ensure the samples were representative of a population. All samples from day 1 to 4 were pooled for sorting and sequencing due to the low frequency of P14 cells in these samples. Enriched samples from the spleen or cell suspensions from lymph nodes were stained with *α*-CD8-PerCP, *α*-CD45.1-APC and fixable lifedead marker to sort live P14 cells (ARIA cell sorter, BD Biosciences).

#### Single-cell RNA sequencing & analysis

Sorted P14 cells from different time-points were washed and resuspended in 0.04% BSA. The single cell sequencing was performed at the Functional Genomics Center Zurich. The cell lysis and RNA capture was performed according to the 10XGenomics protocol (Single Cell 3’ v2 chemistry). The cDNA libraries were generated according to the manufacturer’s protocol (Illumina) and further sequenced (paired-end) with NovaSeq technology (Illumina). The transcripts were mapped with 10Xgenomics CellRanger pipeline (version 2.0.2).

#### Pre-processing & Normalization

Read counts were realigned and sorted for spliced and unspliced counts using the analysis pipeline from velocyto [11]. Contaminating other cell types were removed from the dataset based on outliers in diffusion components. Reads were filtered and normalized according to the Zheng recipe [12] of the scanpy analysis pipeline [23] retaining 5000 highly variable genes. Louvain clustering and UMAP projection were computed using standard parameters, using the first 50 principle components.

#### RNA velocity

RNA velocity uses the relative abundance in reads of un-spliced to spliced mRNA to infer the future state of a particular cell, with a high ratio of un-spliced / spliced mRNA being indicative of recent gene activation, a balanced ratio of un-spliced / spliced mRNA being indicative of gene expression equilibrium, and a low rate of un-spliced / spliced mRNA being indicative for terminating gene expression. Integrating the expression levels of the corresponding genes in neighboring cells allows computing likely transitions between different cellular states in our data set, revealing a vector field demarcating likely transitions.

Scvelo [24] was used to estimate RNA velocity and infer transition probabilities between cells. The transition probabilities were used to construct a Markov process. Inference of RNA velocity relies an a set of assumptions that can be adjusted through several parameters before analysis. The type of pre-processing used, the number of principle components used as well as the neighborhood size for imputation may influence the resulting transition matrix. We conducted an extensive assessment of a many parameter combinations to validate the stability of our RNA velocity analysis. Comparing the equilibrium distributions across 574 parameter sets revealed that the global structure of the transition matrix was quite stable. The parameter set we have chosen for further analysis produced results found in an overwhelming majority of tested sets (Supp. Fig. S5).

We estimated gene moments using neighborhood connectivities using 50 principle components, 30 neighbors with the UMAP method. Velocity was inferred using “stochastic” mode.

#### Cytopath

Cytopath is RNA velocity based lineage inference tool [15]. We applied scvelo’s terminal states routine to compute equilibrium distributions of the forward and backward Markov process, excluding self transitions. Regions with terminal state probability higher than 0.3 were identified and the louvain clusters corresponding to these regions used as start and endpoints. Markov simulations were initialized at the start points and simulated for a maximum of 2000 steps or until they reached the endpoint. We simulated 2000 trajectories from random cell states in the starting region. All simulated paths were aligned to average trajectories from startpoint to each endpoint using dynamic time warping. Neighboring cells (2000 nearest neighbors) were aligned to the trajectories using an alignment score, which was computed based on distance and cosine distance between the cell’s velocity and the direction of the trajectory. Cells that aligned to only one trajectory were assigned to the exhausted or memory-like branch, respectively. Cells at the beginning of the infection that aligned to both trajectories were assigned to the pre-commited branch.

All cells were assigned exhausted, memory-like or pre-commited fate according to their alignment score to the trajectories. Gene expression profiles of 650 genes coding for antibody stainable proteins were then used to predict these labels using L1-penalized Logistic Regression. We used cross validation to identify the optimal L1-penalty that would give a reasonably small number of genes but still good prediction accuracy at C=0.1. The resulting prediction using 12 proteins still classified most cells correctly (accuracy: 0.85). These proteins were then stained on P14 cells from chronic infection at d5 to sort the branches and adoptively transfer them into infection matched hosts.

#### Adoptive transfer experiments

After 5 days of chronic infection, CD8 T cell enriched samples from the spleen or cell suspensions from lymph nodes were stained with *α*-CD8-PerCP, *α*-CD45.1-APC/FITC and *α*-Cxcr6-PE and *α*-Ly108-APC to sort P14 cells into the exhausted, memory-like and pre-commited populations. (ARIA cell sorter, BD Biosciences).

Sorted cells from exhausted (10^6^ cells), memory-like (2×10^5^ cells) and pre-committed (5×10^4^ cells) populations were transferred via intravenous (IV) injection into infection matched hosts infected with either Clone-13 or Clone-13 P14esc mutant. Cells were recovered from from spleens of these mice 12 days post infection prior to phenotypic characterization.

#### Flow cytometry

Surface staining was performed at room temperature for 30 minutes in FACS buffer (2% FCS, 1% EDTA in PBS). LIVE/DEAD™ Fixable Near-IR Dead (Thermo Fisher) was used to discriminate alive from dead cells. Fluorophore-conjugated antibodies used for flow cytometry were purchased from BioLegend (Lucerna Chem AG, Luzern, Switzerland) (*α*-CD45.1 BV711 A20; *α*-CD45.1 APC A20; *α*-CD8 PerCP 53-6.7; *α*-CD8 BV395 53-6.7; *α*-PD-1 PE-BV605 29F.1A12; *α*-Cxcr6 PE SA051D1; *α*-Ly108 APC 330-AJ; *α*-CD8 BV395 53-6.7). Data was acquired LSR II Fortessa using Diva software (BD Biosciences, Allschwil, Switzerland) and analyzed in FlowJo (BD Biosciences, Allschwil, Switzerland). Gating and plotting was done using FlowJo (BD Bioscience, Allschwil, Switzerland).

## Aknowledgement

We thank Prof. Roman Spörri for providing the Nr4a1-GFP mice. We thank the Pinschewer laboratory through the European Virus Archive (University of Basel) for providing the Clone-13 P14 escape mutant viral strain. We thank Franziska Wagen and Nathalie Oetiker for great technical support. We are grateful for the constructive input of the members of the Claassen, Oxenius, Joller and Sallusto Group during discussions and group meetings. Funding: This work was support by the ETH Zürich (grant no. ETH-39 14-2 to MC and AO) and Novartis.

## Contributions

D.C., M.C., A.O. designed the experiments; D.C., I.S. carried out the experiments, D.C., R.G., M.C., A.O. analyzed the experiments; D.C., M.C., A.O. wrote the manuscript.

## Data Availability

Sequencing Data is available on request from the authors.

