## Supplementary Figures for "Fate trajectories of CD8^+^ T cells in chronic LCMV infection"

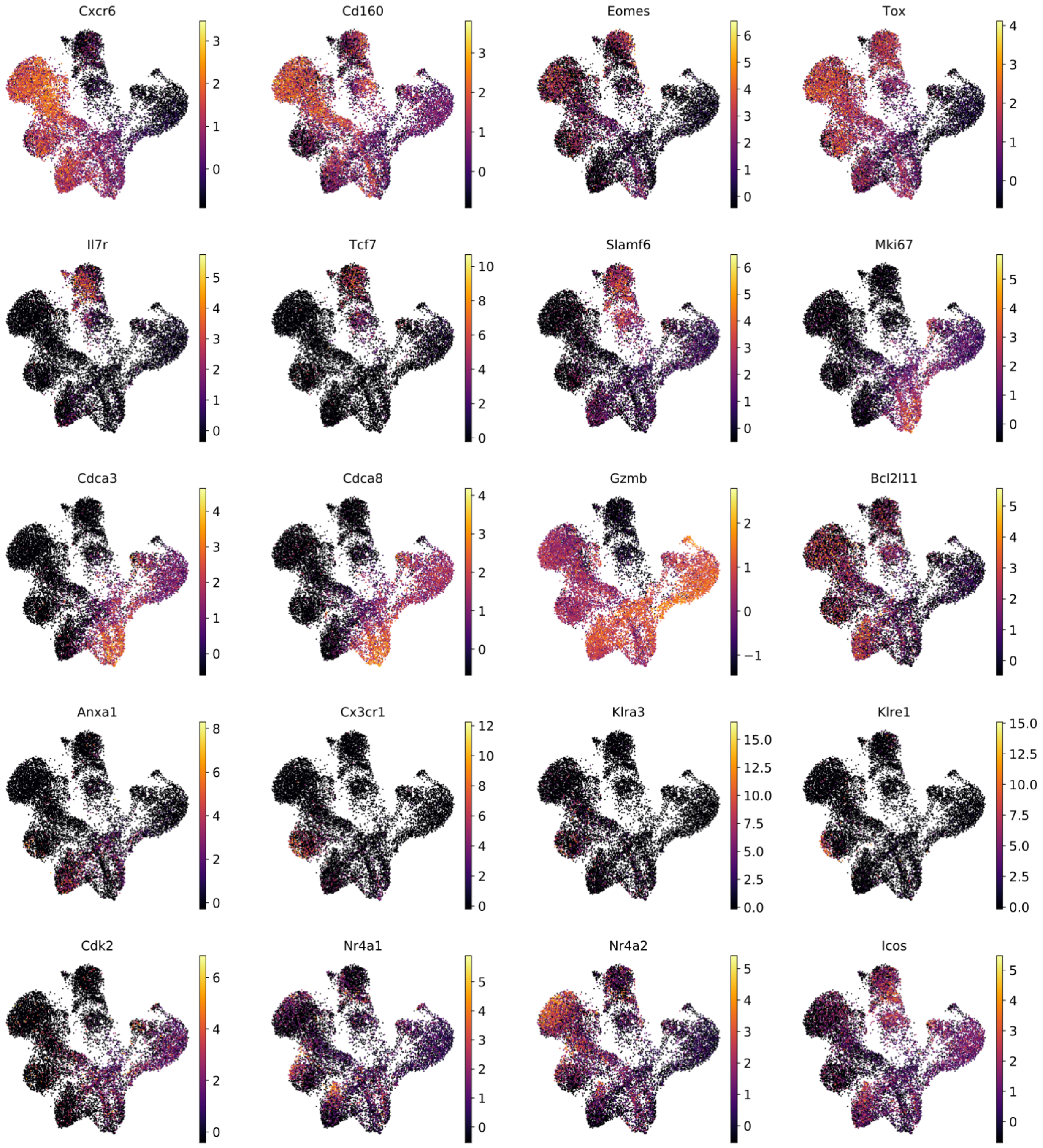


Figure S1: distribution of relevant genes

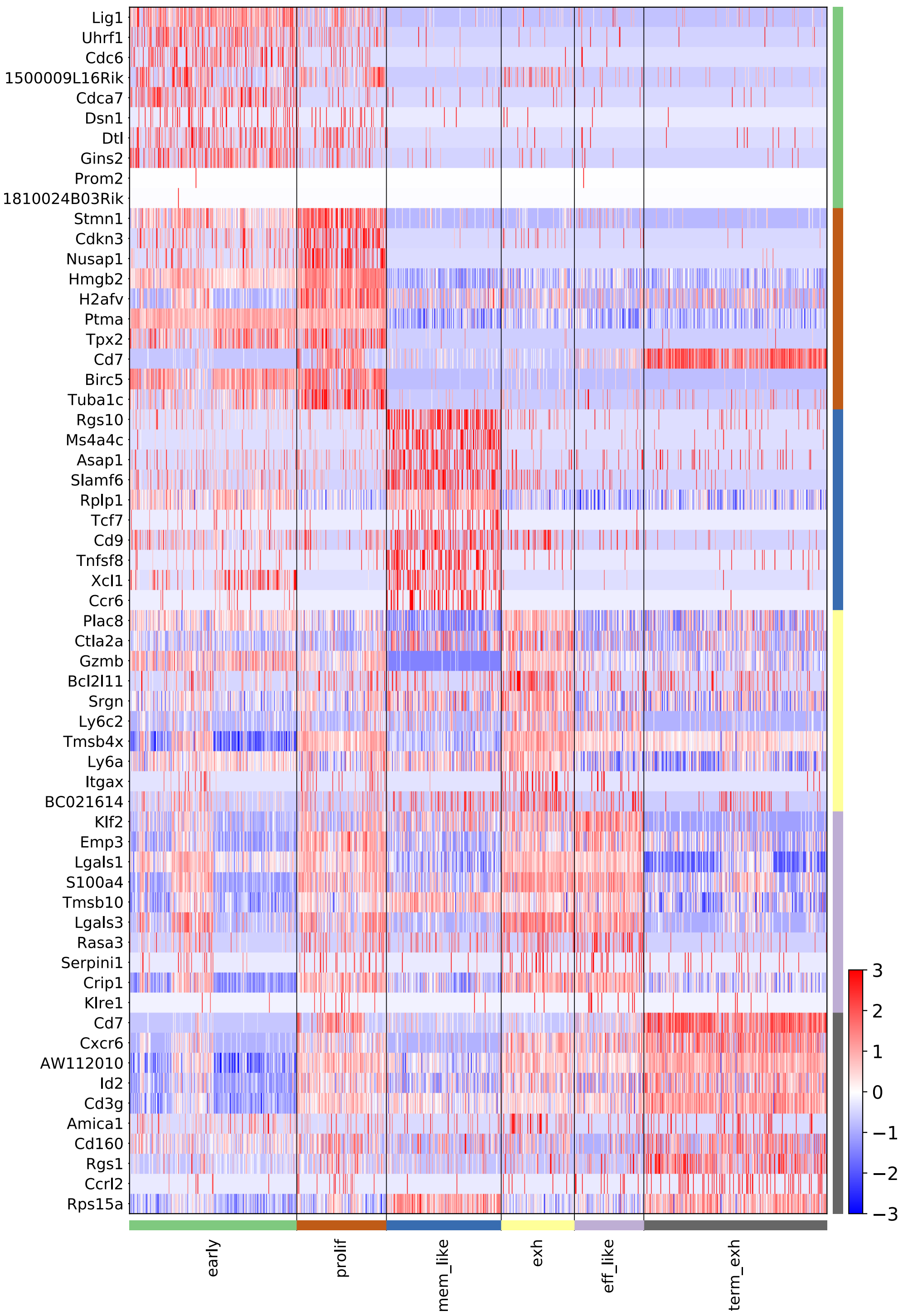


Figure S2: Top 10 differentially expressed genes between relevant populations

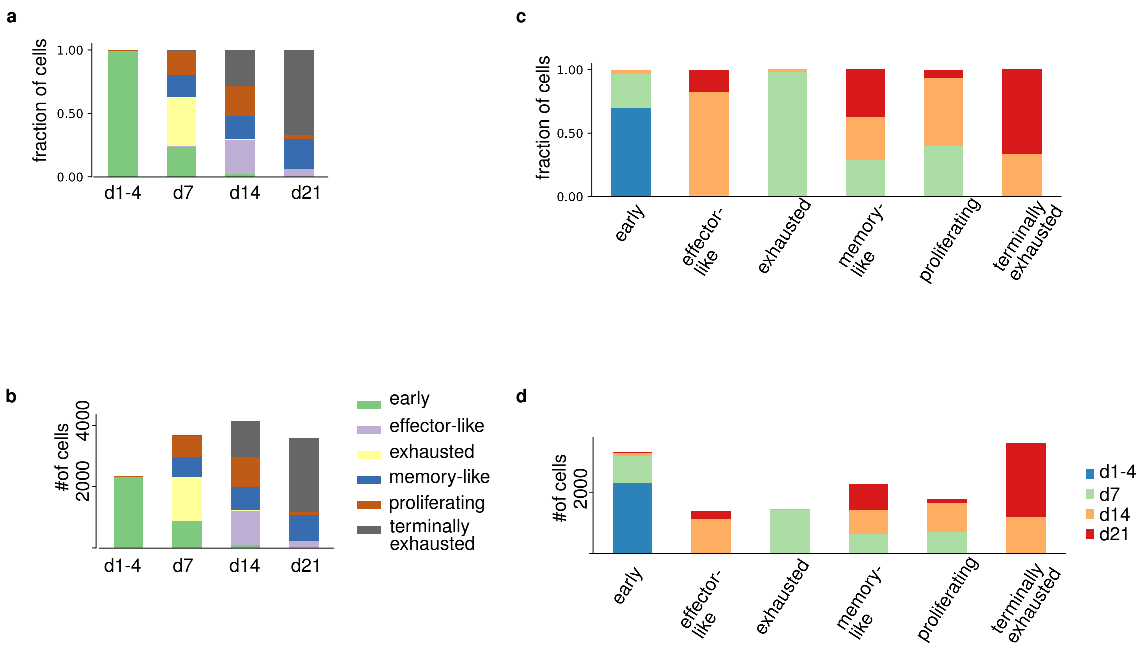


Figure S3: Composition of phenotyic groups with respect to samples. (a) fractional composition per sample time-point (b) cell numbers per group per sample time-point (c) fractional composition per phen. group (d) cell numbers per group per sample time-point

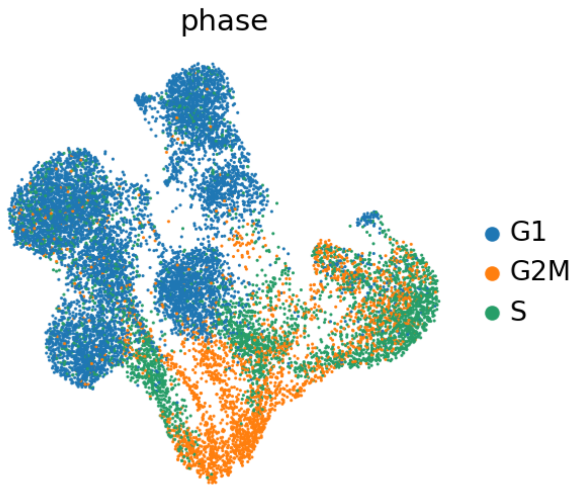


Figure S4: cell cycle scoring confirmed strong G2M activity in the proliferating group

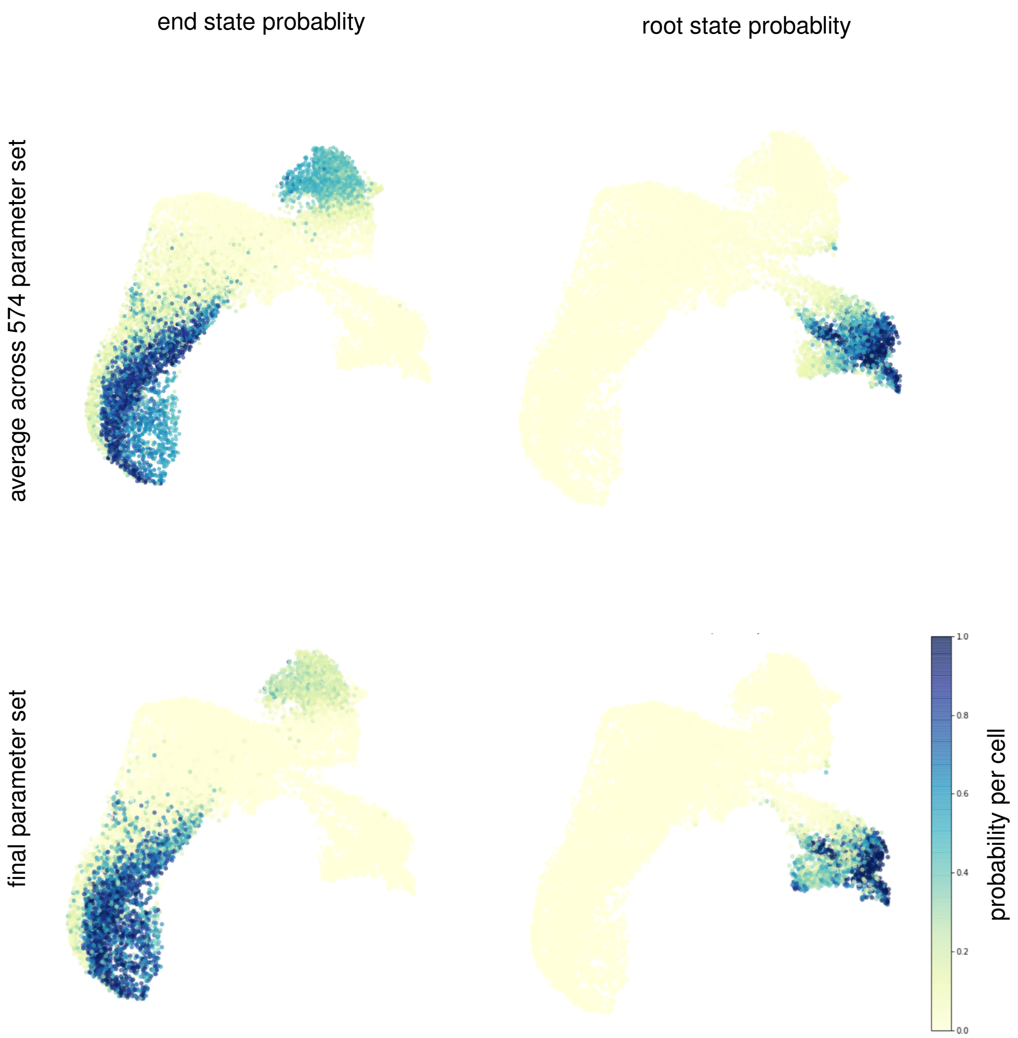


Figure S5: mean end and root point probability across many parameters

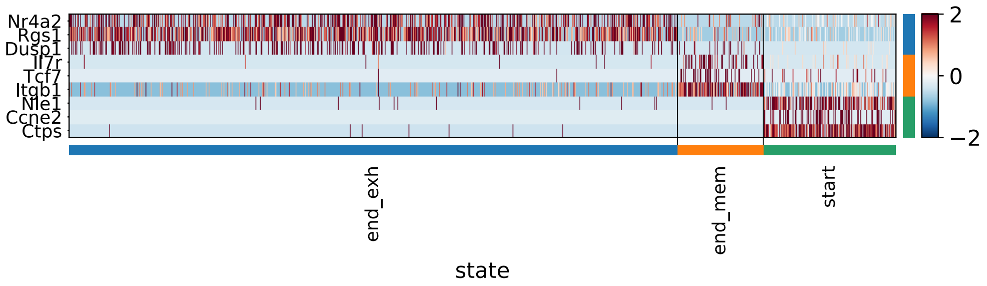


Figure S6: differentially expressed genes in root and end states

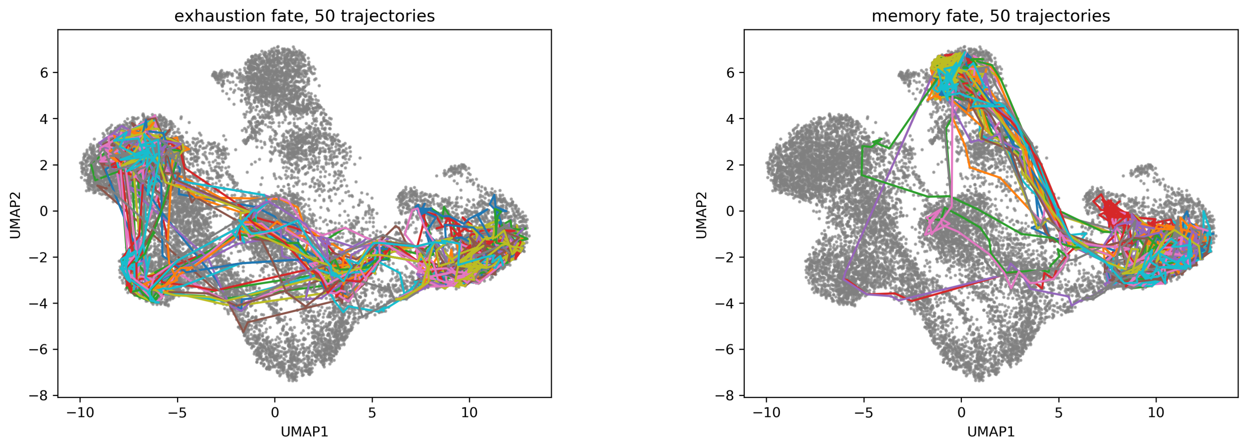


Figure S7: Simulated trajectories from starting region to exhaustion endpoint

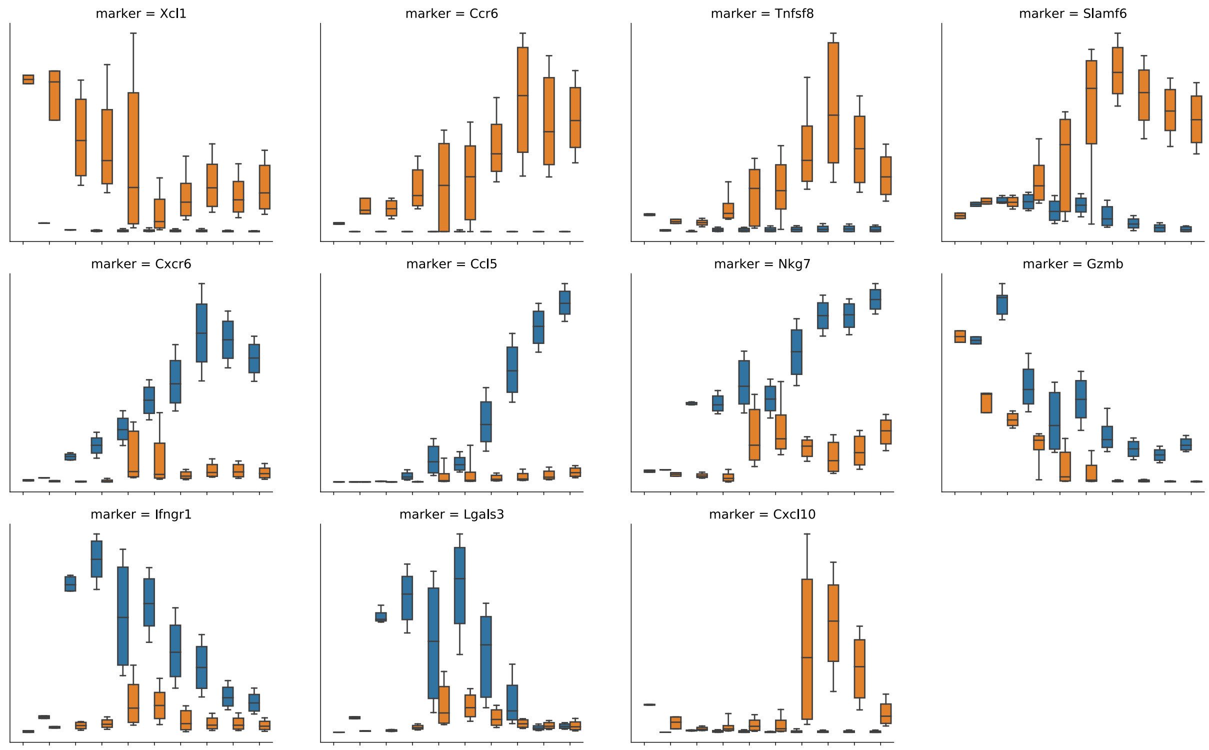


Figure S8: selection of differentially expressed genes between memory-like and exhausted trajectories in order of their significance score.

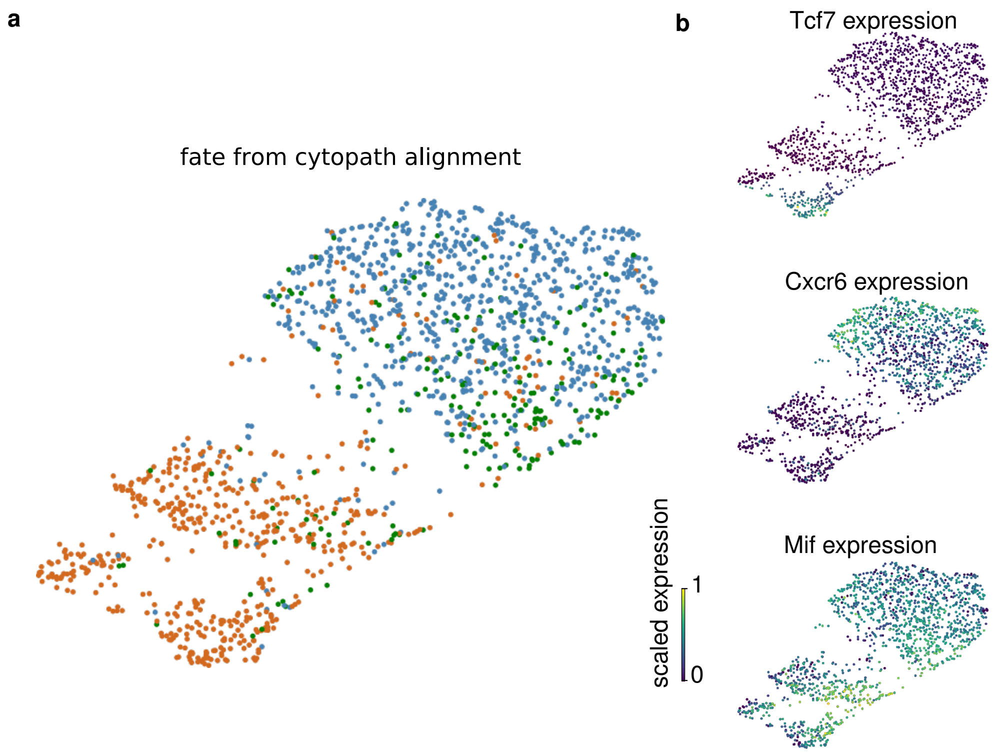


Figure S9: Using only identified marker genes allowed good precition of the assigned branch fate.
(a) Three branches in umap projection.
(b) Branch specific gene expression.
